# Species Diversity and Mitochondrial DNA Analysis of Sponges (Porifera, Demospongiae) in an Anchialine Cave on the Yucatan Peninsula

**DOI:** 10.1101/2024.10.31.621400

**Authors:** Pablo Suárez-Moo, Adán J. Rivera-Domínguez, Norma A. Márquez-Velázquez, Alejandra Prieto-Davó

## Abstract

The anchialine ecosystem in the southeastern Gulf of Mexico has an unexplored fauna, among the most understudied groups is Porifera, molecular approaches to investigate their biology and evolution remain largely short. To broaden these studies, sponge individuals collected in the anchialine cave “Xcalac” in the Mexican state of Quintana-Roo were analyzed using 18S rRNA sequences from sponge metagenomes. The three individuals studied belong to the *Cinachyrella, Xestospongia, and Suberites* genera. The mitochondrial (mt) genomes of *Cinachyrella* sp. n. and *Xestospongia* sp. n. were 18,493 bp to 19,604 bp in length, containing 14 protein-coding genes, 2 rRNA genes (rrnS and rrnL), and 23–25 tRNA genes, respectively. The phylogenomic analysis showed that *Cinachyrella* sp. n. had the same gene arrangement as the members of its subclade, including sponge species of the *Cinachyrella* and *Geodia* genera. The mt genome of *Xestospongia* sp. n. contained the same gene arrangement as found in other sponges of the same genus and differed from other genera such as *Petrosia* and *Haliclona* by an tRNA (tyrosine, Y1). Variation in mitochondrial genomes (size, gene content, and gene order) was observed when comparing sampled sponge species from the class Demospongiae to the class Homoscleromorpha. This is the first record of *Cinachyrella* sp. n and *Suberites sp. n.* in an anchialine cave on the southeastern Yucatan peninsula, and the first report of the mitochondrial genome analysis of *Cinachyrella* sp. n. and *Xestospongia* sp. n., contributing to a better understanding of the diversity and phylogeny of sponges in this ecosystem.

## Introduction

The Yucatan peninsula is a limestone platform with a subterranean aquifer ecosystem that has unique geological and chemical conditions (Bauer-Gottwein et al. 2011). These conditions have led to the formation of an underwater anchialine cave system approximately 330 km in length (Calderón-Gutiérrez et al. 2017). This ecosystem can exhibit extensive horizontal branching, connections to the sea through conduits, and increasing depth away from the coastline (Benítez et al. 2019). These caves are characterized by light limitation, pressure and temperature variations, varying levels of dissolved organic matter, and fluctuating oxygen availability (Yañez-Mendoza et al. 2007, Romero 2019). Such physicochemical conditions may result in high species richness and endemism rates reported for these ecosystems (Yañez-Mendoza et al. 2007, Calderón-Gutiérrez et al. 2017, Gómez and Calderón-Gutiérrez 2020). Previous investigations of the fauna in the anchialine caves of the region have focused on phytoplankton and tychoplankton communities (Sánchez et al. 2002), Crustacea (Alvarez et al. 2015, Benítez et al. 2019, Álvarez et al. 2023), Echinodermata (Mejía-Ortíz et al. 2007, Bribiesca-Contreras et al. 2019), and other groups such as Annelida and Mollusca (Álvarez et al. 2023). A particular group of interest is the phylum Porifera (sponges), which can enhance primary production, participate in biogeochemical cycles, and provide microhabitats for a wide range of organisms (Koukouras et al. 1996; Pérez-Botello and Simões 2021). A high number of sponge species have been reported in the anchialine caves of Cozumel island in the Yucatan peninsula (Calderón-Gutiérrez et al. 2017, 2018, Gómez and Calderón-Gutiérrez 2020), in coral reefs in the Caribbean Sea and the Gulf of Mexico (Ugalde et al. 2021, Pérez-Botello et al. 2023), and in mangrove roots from the southern Gulf of Mexico (Castellanos-Pérez et al. 2020). Reports of sponges in the anchialine caves of the mainland Yucatan peninsula are scarce, and there is only one study that reported the microbial communities and biotechnological potential of bacteria isolated from a sponge belonging to the genus *Xestospongia*, collected in the anchialine cave of Xcalak, Quintana Roo, México (Suárez-Moo et al. 2024). In contrast, marine sponges in the tropics have been broadly studied. including topics such as sponge ecology (Wulff 2006, Araya-Vargas et al. 2020), new species’ description(Shen et al. 2022, Shilov et al. 2023, Díaz et al. 2024), diversity of the microbial communities associated to their tissues (Busch et al. 2022), metabolic potential of their microbial communities (Lesser et al. 2022), and biotechnological applications of their isolated bacteria (Jayatilake et al. 1996, Su et al. 2014, Santos et al. 2015).

A common molecular marker used to determine the phylogenetic relationships among different animals is the mitochondrial DNA (mtDNA). MtDNA-based phylogenetic trees are considered valuable tools for understanding evolutionary history within and between most groups of metazoans (Feng et al. 2023). Mitochondrial genome analysis has been used to understanding the phylogeny (Zhang et al. 2016, Taboada et al. 2018), mtDNA evolution (Wang and Lavrov 2008) and divergence time (Plese et al. 2021) of sponges in different environments.

The lack of information about the biodiversity and genomic characterization of the fauna from anchialine caves in the Yucatan peninsula is concerning due to their important ecological roles in the karst ecosystem and to the potential changes in their diversity in the current context of climate change. Additionally, the region is experiencing increasing human activity impacts due to the expansion of tourism and urban development.

In this study, we describe three sponge species (*Cinachyrella* sp. n., *Xestospongia* sp. n., and *Suberites* sp. n.) from an anchialine cave on the coastal area in the Yucatan peninsula, including new records for two of these species. We also provide the annotated mitochondrial genomes of *Cinachyrella* sp. n. and *Xestospongia* sp. n., compare their gene arrangements with those of closely related species, and discuss the phylogenetic framework for these sponges using the nucleotide of their respective amino acid sequences

## Material and methods

### Sample collection and preservation

One specimen from each sponge species belonging to the class Demospongiae was collected at a depth of approximately 12 meters inside an anchialine cave (“Cayo Judio”, Xcalak) from the underground karst aquifer sinkhole in the Yucatan Peninsula in September 2020, (Figure 1; for details see Suarez-Moo et al. (Suárez-Moo et al. 2024)). The sponges were photographed, removed from the nodule with a scalpel, preserved in RNALater, and transported to the lab at 4 °C and immediately stored at −20 °C until DNA extraction. The sponges’ samples were identified using a traditional taxonomic approach based on the tissues and spicules (Gómez and Calderón-Gutiérrez 2020). The samples were collected under the Secretaría de Agricultura, Ganadería, Desarrollo Rural, Pesca y Alimentación (SAGARPA) official permit number PPF/DGOPA-062/21.

**Figure 1.**
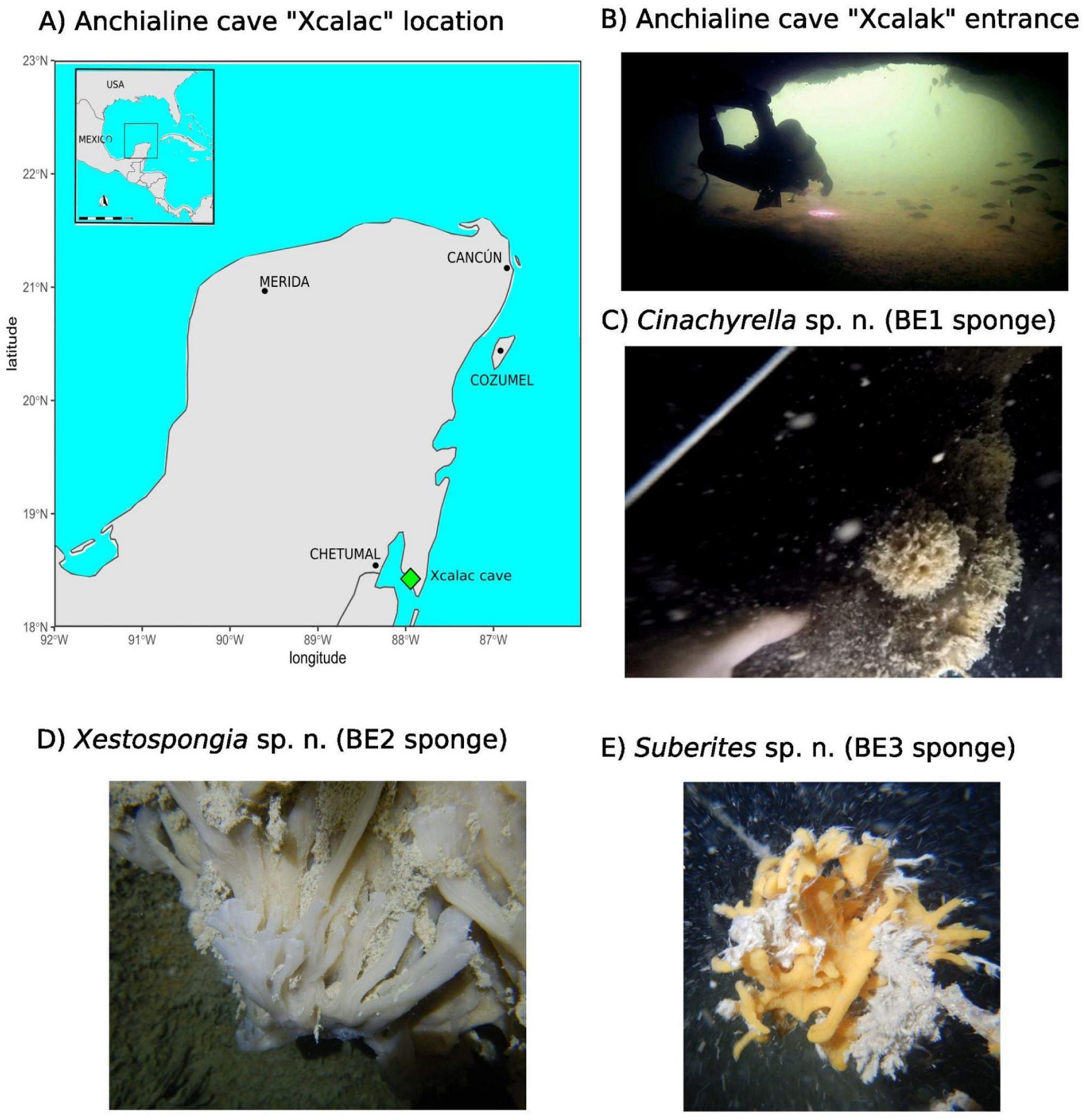
Collection site location and sponge specimens. (A) Study area: anchialine cave “Xcalak” (lime diamond), located in the southeast Gulf of Mexico. (B) Photograph of the cave entrance. (C) BE1 sponge. (D) BE2 sponge. (E) BE3 sponge. All sponges were collected in the anchialine cave “Xcalak.”

### DNA extraction and illumina sequencing

Genomic DNA from approximately 1 cm³ of the sponge tissues was extracted in triplicate using a ZymoBIOMICS Miniprep D4300 Kit (Zymo Research, Irvine, USA). DNA from the triplicates was pooled, and its purity and concentration were analyzed using a spectrophotometer (Nanodrop 2000, Thermo Fisher Scientific, Waltham, USA) and visualized by agarose gel electrophoresis. The DNA was stored at –20 °C for further processing. The extracted DNA was subjected to quality control (Qubit Fluorometer), library preparation, and shotgun sequencing by Novogene (Davis, California, USA), on a NovaSeq. 6000 platform (Illumina).

### Taxonomic assignation

The quality of the raw reads was checked with FastQC version 0.11.2 (Andrews, 2010) and multiQC (Ewels et al. 2016). Low-quality bases (per base sequence quality <33) were removed with Trimmomatic version 0.39 (Bolger et al. 2014). High-quality reads (encompassing prokaryotic and eukaryotic reads) were aligned against a nonredundant version of the SILVA database v138 (Quast et al., 2013) with an E-value <10−5. Sequences matching this database were considered potential rRNA gene fragments, and they were aligned against Eukarya hidden Markov models (HMMs) using SSU-ALIGN v. 0.1.1 (Nawrocki, 2009) to identify true sequences. The Eukarya rRNA reads were assembled in rRNA contigs with metaSPAdes version v. 3.13.0 (Nurk et al. 2017). A mapping of the Eukarya rRNA reads against the different Eukaryotic rRNA contigs from the three sponges samples was performed using the software Bowtie 2 (Langmead and Salzberg 2012). To get an estimate of the mean coverage of each Eukaryotic rRNA contig, the following formula was used: (Depth of coverage) × (Number of bases on contig)/ (Length of the contig). The frequency of the mean coverage of the Eukaryotic rRNA contigs detected in each sponge library was used in an alluvial diagram with the package ggplot2 (Wickham 2016).

The Eukaryotic rRNA contigs assembled were classified using a BLASTN (Altschul et al., 1997) against the SILVA taxonomy database (Quast et al., 2013) and NCBI database. The contigs that matched the 18S rRNA sequences of the Porifera phylum were extracted, and reference sequences downloaded from the GenBank database (Table S1) were aligned using Mafft v.7.305b (Katoh and Standley 2013) and adjusted with trimAl v1.4.rev15 (Capella-Gutiérrez et al. 2009). Phylogenetic estimation was performed using IQ-TREE (Nguyen et al. 2015) with 10,000 ultrafast bootstraps and the -m MFP option to find the best model that fitted our data. The phylogenetic tree was visualized with iTOL (Letunic and Bork 2019). A parallel study of the sponge microbiome was conducted in our lab, therefore the prokaryotic reads and contigs of these metagenomes were not analyzed in this study. The nucleotide sequences of the 18S rRNA contigs assembled in this study were deposited in the NCBI under the accession number PQ492272-PQ492274

### Mitochondrial genome assembly and annotation

The high-quality reads (encompassing prokaryotic and eukaryotic reads) were used to map reads back to the reference mitochondrial genomes belonging to genera identified in the 18S rRNA analysis (Table S2) using Bowtie 2 (Langmead and Salzberg 2012). SAMtools v.1.9 (Li and Durbin 2009) was used to convert files to binary format for further downstream analyses. These mapped reads were used for the mitogenomes assembly using metaSPAdes version v. 3.13.0 (Nurk et al. 2017). The mitogenomes were annotated using web server MITOS2 (Bernt et al. 2013) with the Ascidian NCBI code for translation, and subsequently manually curated and visualized with Geneious Prime 2024.0.5 (Biomatters Ltd, Auckland, New Zealand). Transfer RNA genes were identified using the tRNAscan-SE program (Lowe and Eddy 1996) and MITOS2 (Bernt et al. 2013).

For the phylogenomic analysis, the nucleotide sequences of all protein-coding genes, along with rrnS and rrnL sequences from the BE1 and BE2 sponge mitogenomes and reference mitogenomes downloaded from the GenBank database (Table S2) were concatenated using custom bioinformatic commands and aligned using Mafft v.7.305b (Katoh and Standley 2013). The length of the alignment was adjusted with trimAl v1.4.rev15 (Capella-Gutiérrez et al. 2009). The phylogenomic tree was performed using IQ-TREE (Nguyen et al. 2015) with 10,000 ultrafast bootstraps and the -m MFP option to find the best model that fitted to our data. The phylogenomic tree was visualized with iTOL (Letunic and Bork 2019).

## Results

### Taxonomic assignation of the three sponges species from the “Xcalak” anchialine cave

The morphology analysis indicated that the samples BE1, BE2, and BE3 were sponges from the orders Tetractinellida, Haplosclerida, and Suberitida respectively. Using standard identification keys based on spicule analysis, *Cinachyrella* was identified as the candidate genus for sponge BE1, *Haliclona* or *Xestospongia* were potential candidates for BE2, while the family Suberitidae was identified as the candidate for the spongeBE3 in this study. The taxonomic identification of the three sponges was confirmed through the taxonomic assignment of rRNA contigs assembled from the metagenomic shotgun data.

In the taxonomic assignment of the rRNA contigs for the BE1 sponge, five contigs were assembled. The longest contig with the most coverage, (1704 bp) belonged to the genus *Cinachyrella* (order Tetractinellida, phylum Porifera) (Figure 2A and Table S3). However, other organisms, including sponges from the genus *Suberites* (with a low-abundance, and short contig) and the genera *Erinaceusyllis* and *Syllis* (order Phyllodocida, phylum Annelida), were also associated with the BE1 sponge (Figure 2A and Table S3). Thirteen contigs were assembled for the sponge BE2. Three of them were assigned to the genus *Xestospongia* (order Haplosclerida, phylum Porifera), including one contig with the highest coverage and a length of 2029 bp (Figure 2A and Table S3). A short contig (171 bp) belonging to the genus *Suberites* (order Suberitida, phylum Porifera) was also assembled from this sample (Figure 2A and Table S3). Another long contig (1830 bp) was assigned to the genus *Diadumene* (order Actiniaria, phylum Cnidaria) (Figure 2A and Table S3). The three contigs assembled in the sponge BE3 were assigned to the genus *Suberites* (order Suberitidae, phylum Porifera), with the longest contig measuring 2048 bp (Figure 2A and Table S3).

**Figure 2.**
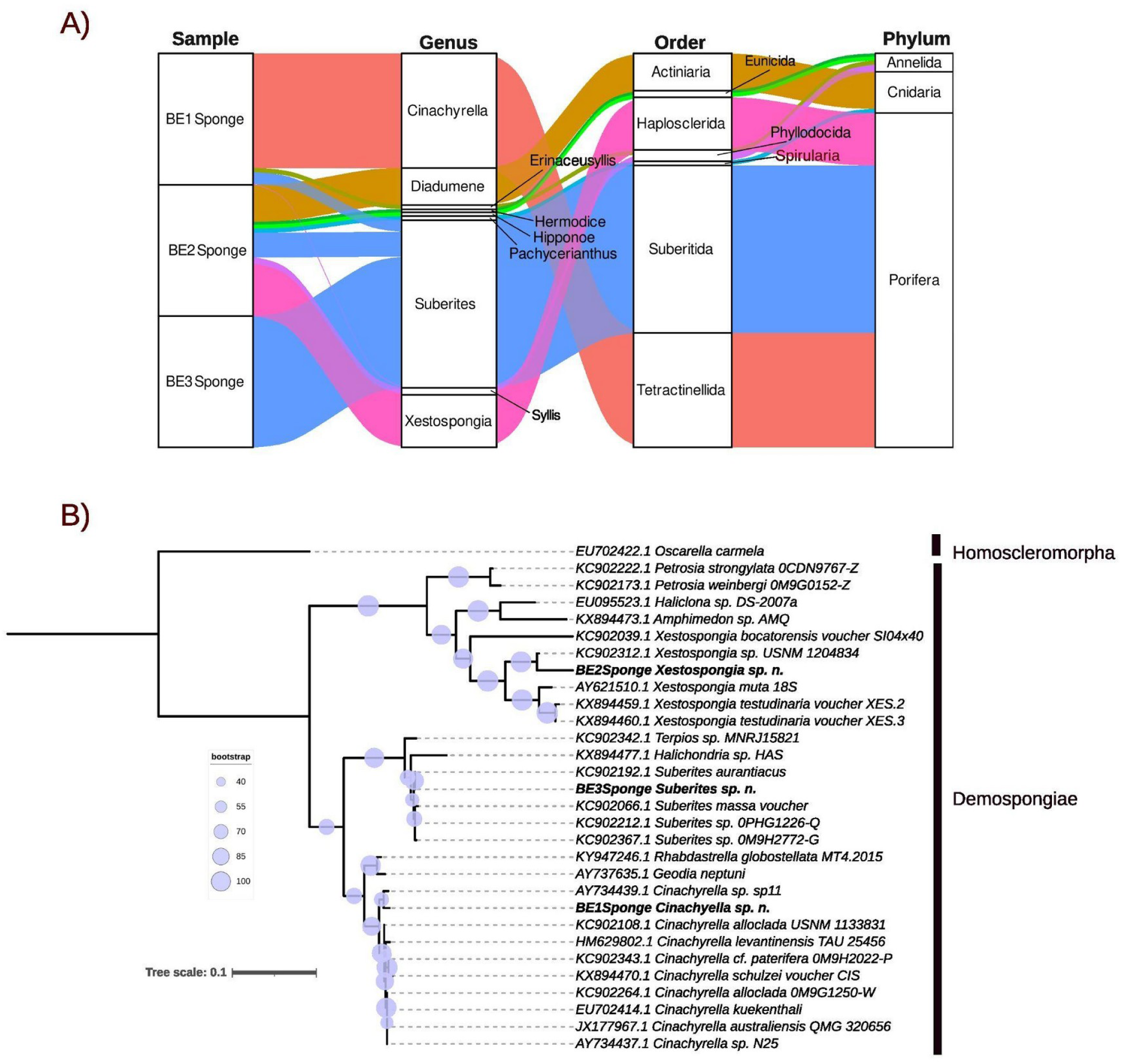
Taxonomic identification and phylogenetic analysis of three sponges species collected in the anchialine cave “Xcalak”. (A) The rRNA contigs were taxonomically assigned to genus, order, and phylum levels, represented by different colors. (B) 18S rRNA contigs assembled from the complete metagenomic data were used, along with reference sequences, in a phylogenetic analysis. The circles and the width of each node are Bayesian posterior probabilities.

Using a BLASTn against the NCBI database, the 18S rRNA sequence of sponge BE1 had a 99.6% identity with *Cinachyrella* sp. Sp11 (AY734439.1) and 98.8% with *Cinachyrella schulzei* (KX894470.1). The BE2 sponge had 98% identity with *Xestospongia sp.* USNM 1204834 (KC902312.1) and 93% with *X. muta* (AY621510.1). The BE3 sponge had a 100% identity with *Suberites aurantiacus* voucher MNRJ15701 (KC902192.1) and 99.65% with *S. massa* voucher BELUM:Mc4528 (KC902066.1).

### Phylogeny and mitochondrial genome analysis

The 18S rRNA phylogenetic analysis using reference organisms confirmed taxonomic affiliations and showed that the BE1 sponge clustered with species of the genus *Cinachyrella*, BE2 clustered with those from *Xestospongia*, and the BE3 sponge with *Suberites* (Figure 2B). Evolutionary speaking, sponge BE1 was more closely related to sponge BE3 than to the BE2 sponge. Therfore, based on the taxonomic assignment of the rRNA contigs and 18S rRNA phylogenetic analyses, the sponges BE1, BE2, and BE3 were conservatively identified as *Cinachyrella* sp. n., *Xestospongia* sp. n., and *Suberites* sp. n., respectively.

In this study, only the mt genomes of *Cinachyrella* sp. n. and *Xestospongia* sp. n. were used, as *Suberites* sp. n. sequences had a low-quality assembly characterized by a small number of short contigs, and poor gene characterization. The mt genome of *Cinachyrella sp. n.* consists of 18,493 bp (Figure 3A) and includes 23 tRNA genes, 2 rRNA (rrnS and rrnL), and 14 protein coding genes. These include three subunits of the ATP synthase (atp6, atp8, atp9), seven NADH dehydrogenase subunits (nad1, nad2, nad3, nad4, nad4l, nad5, nad6), one apocytochrome b (cob), and three Cytochrome C oxidase subunits (cox1, cox2, cox3) (Figure 3A). All genes are positioned on the heavy strand and are transcribed counter-clockwise. The mt genome of *Xestospongia sp.* consists of 19,604 bp (Figure 3B) and contains 25 tRNA genes, 2 rRNA (rrnS and rrnL), and 14 protein coding genes, including three subunits of the ATP synthase (atp6, atp8, atp9), seven NADH dehydrogenase subunits (nad1, nad2, nad3, nad4, nad4l, nad5, nad6), one apocytochrome b (cob), and three Cytochrome C oxidase subunits (cox1, cox2, cox3)(Figure 3B). All genes are positioned on the heavy strand and are transcribed clockwise.

**Figure. 3.**
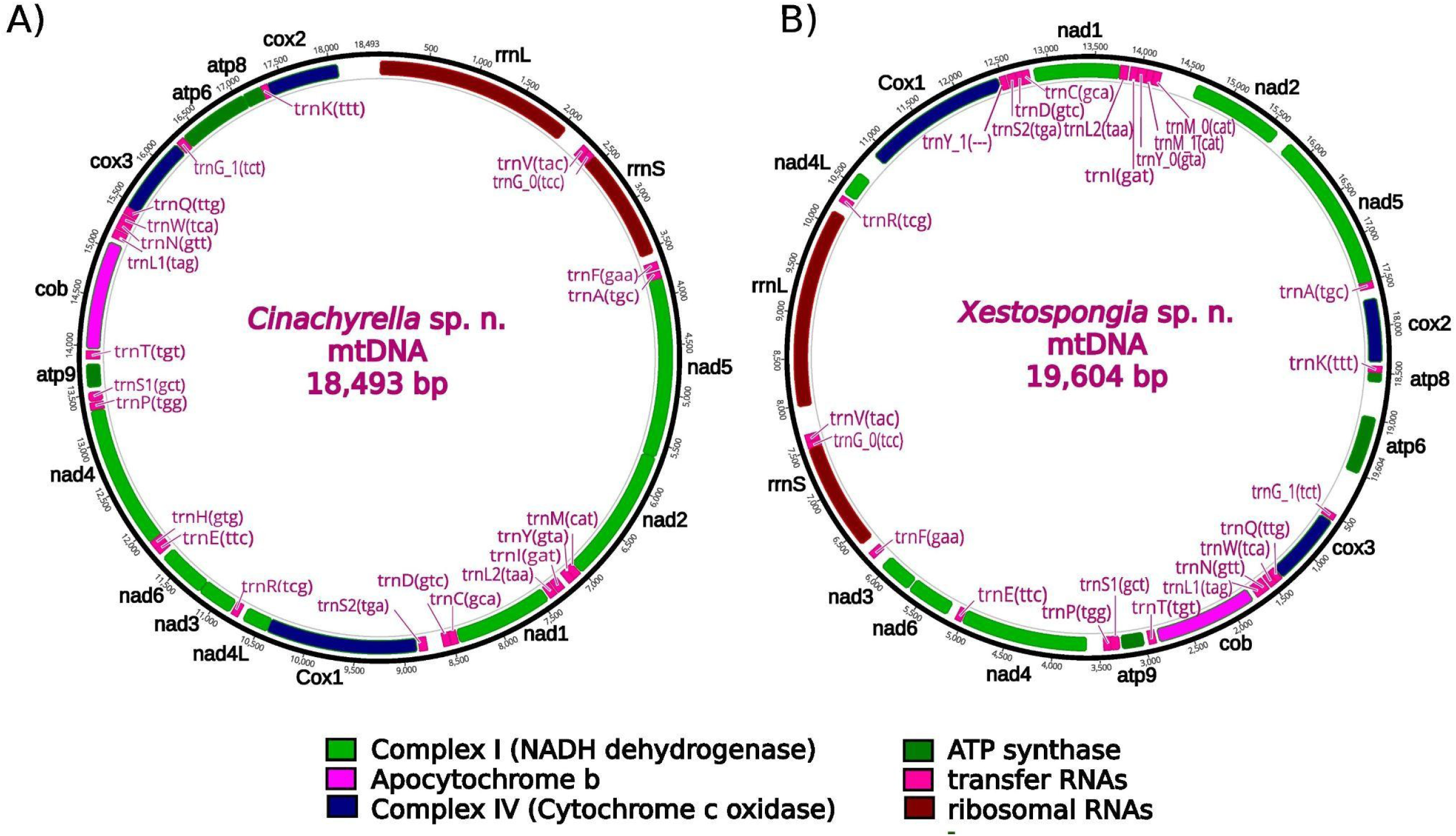
Genetic map of *Cinachyrella* sp. n. and *Xestospongia* sp. n. circular mitochondrial genomes. All genes are positioned on the heavy strand and are transcribed (A) counterclockwise for *Cinachyrella* sp. n. and (B) clockwise for *Xestospongia* sp. n.

The phylogenomic tree of the concatenated nucleotide alignment of all protein-coding gene sequences, along with rrnS and rrnL shows two clades (Figure 4A): Clade (A) includes the three sponge species belonging to the class Homoscleromorpha (groups G1 and G2), while clade B contains two subclades. The first subclade includes *Plenaster craigi* (group G3) and species from the genera *Geodia* and *Cinachyrella*, including sponge BE1 *Cinachyrrella* sp. n. (group G4). The second subclade includes species from the genera *Petrosia* and *Haliclona* (group G5) and the genus *Xestospongia* (group G6) including sponge BE2 *Xestospongia* sp. n.

**Fig. 4.**
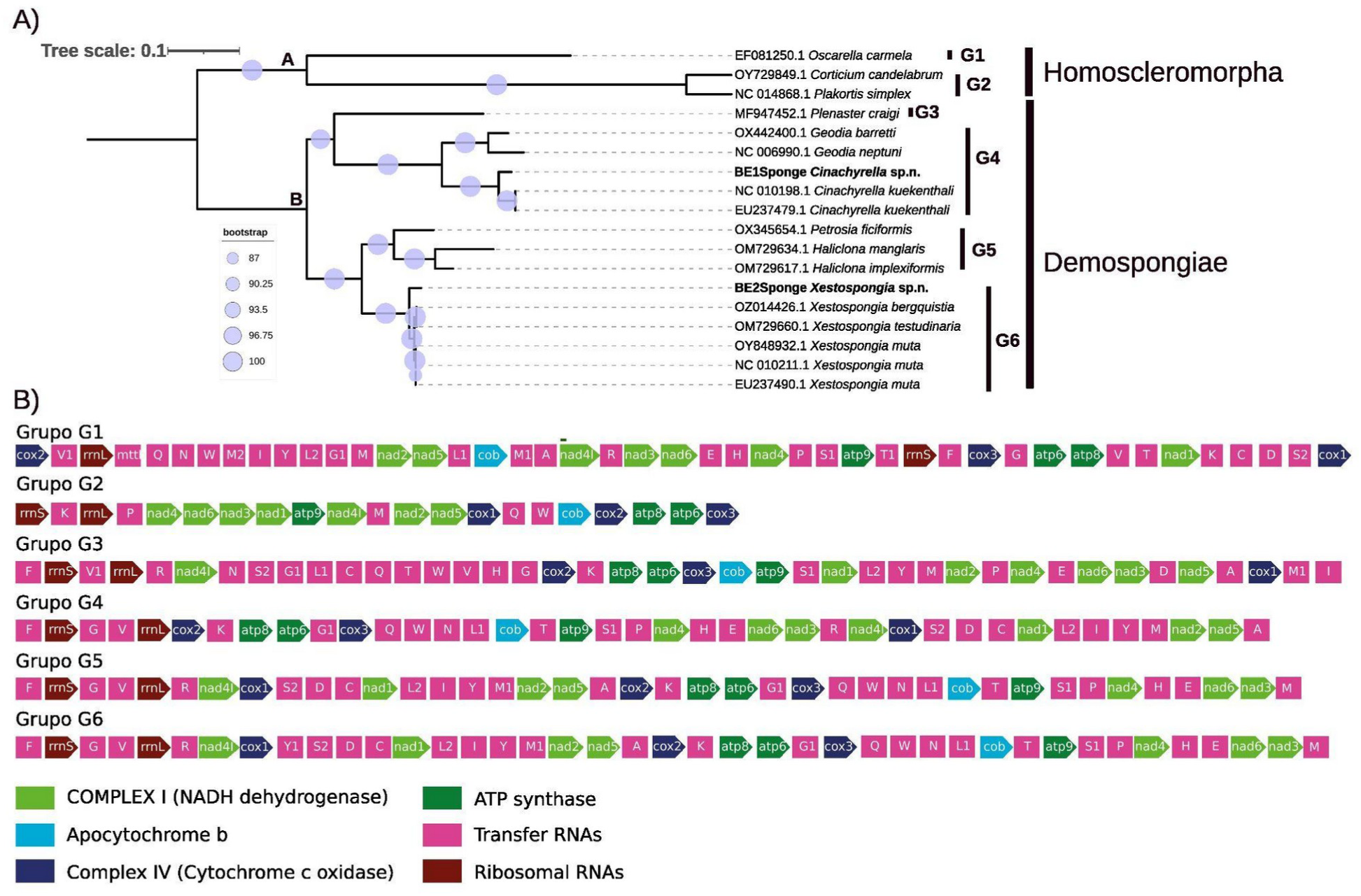
Phylogenomic analysis and mitochondrial gene arrangements between *Cinachyrella* sp. n. and *Xestospongia* sp. n., demosponge species, and outgroup heteroscleromorph. A) Maximum likelihood phylogenetic tree using a concatenated nucleotide alignment of all protein coding gene sequences along with those of rrnS and rrnL. Circles at bases of nodes represent bootstrap support nodes. B) Mitochondrial genome gene order, the complex I (Lime), apocytochrome b (fuchsia), complex IV (Navy), ATP synthase (Green), tRNAs (pink), rRNA genes (Maroon) are shown.

The mt genome of sponge BE1 *Cinachyrella sp. n.* exhibits the same genome organization and gene arrangement as other sponges from the same genus but also from the genus *Geodia* (group G4, Figure 4B). In contrast, sponge BE2 *Xestospongia* sp. n. shows the same genome organization and gene arrangement as found in other sponges of the same genus and lacks a tRNA (tyrosine, Y1) found in other genera such as *Petrosia* and *Haliclona* (group G5 vs group G6, Figure 4B). Groups G4 and G6, to which *Cinachyrella* sp. n. and *Xestospongia* sp. n belong, respectively, differ in gene order and the former lacks the trnM gene found in the latter.

Variation in the size, gene content, and gene order was observed between mitochondrial genomes of sponge species from the classes Demospongiae and Homoscleromorpha (Figure 4B). *Oscarella carmella* (group G1, Homoscleromorpha) has the highest number of tRNAs, with 28, while *Corticium candelabrum* and *Plakortis simples* (group G2, Homoscleromorpha) have the lowest number, with 5 tRNAs (Figure 4B). In contrast, species belonging to Demospongiae (groups G3–G6) have 14 protein-coding genes and 2 rRNAs (rrnS and rrnL), with a range of 22–25 tRNAs (Figure 4B).

## Discussion

Our study represents the first record of the *Cinachyrella sp. n.* and *Suberites* sp. n. in the anchialine system of the Yucatan peninsula, and following the report of *Cinachyrella* sp. n. and *Xestospongia sp.* n. (Suárez-Moo et al. 2024), this is the first mitochondrial genome analysis for both species. Most studies on the characterization of sponges in Mexico’s anchialine caves have been conducted in Cozumel Island (Calderón-Gutiérrez et al. 2017, 2018, Gómez and Calderón-Gutiérrez 2020), highlighting the need to continue exploring the species diversity and the distribution of sponges in these unusual cave systems from the mainland of the Yucatan Peninsula.

The phylum Porifera is highly diverse in anchialine caves such as “La Quebrada” and “El Aerolito”, located in Cozumel Island, with 15 and 29 sponges species reported, respectively, (Table S4). The Xcalak cave only had three sponge genera at the moment it was sampled: *Cinachyrella* sp. n., *Xestospongia* sp. n., and *Suberites* sp. n. with the last two of these being new to the anchialine system, since *Cinachyrella kuekenthali*) was previously reported at “La Quebrada” (Calderón-Gutiérrez et al. 2018; Gómez-Gutiérrez and Calderón-Gutiérrez 2020). In total, 161 sponge species have been reported from Mexican waters in the Gulf of Mexico (i.e. not from a karstik anchialine system) (Ugalde et al. 2021), including species from the genera *Cinachyrella* (*C. alloclada*, *C. apion*, and *C. kuekenthali*), *Xestospongia* (*X. arenosa* and *X. muta*), and *Suberites* (*S. aurantiacus*) (Pérez-Botello and Simões 2021; Ugalde et al. 2021). Differences in the number of sponge species between anchialine caves from Cozumel island and the mainland of the Yucatan Peninsula could be related to variation in cave morphological characteristics. The caves “La Quebrada” and “El Aerolito” are significantly longer, with lengths of 9.2 km and 18 km, respectively (Calderón-Gutiérrez et al. 2017, 2018; Gómez-Gutiérrez and Calderón-Gutiérrez 2020), whereas the length of Xcalak cave has been estimated to be only ∼30 m (divers’ personal observations). Another important difference is that both Cozumel island caves have entrances connected to the Caribbean Sea and coastal sinkholes locally called “cenotes” (Calderón-Gutiérrez et al. 2017, 2018; Gómez-Gutiérrez and Calderón-Gutiérrez 2020), while the cave at Xcalak has a single entrance, a coastal sinkhole known as “Cayo Judío”. Therefore, the anchialine caves could have an energy input to the anchialine system by a mangrove in the main entrance (El Aerolito cave), and the direct connection with the sea leading to an exceptional biodiversity (Calderón-Gutiérrez et al. 2017). On the other hand, this connection to the Caribbean Sea may enhance species diversity in ‘La Quebrada’ and ‘El Aerolito’ caves by transporting marine larvae, including those from deep-sea species, into the anchialine caves via upwelling events (Harmelin & Vacelet 1997) or passively through sea-level changes (Hart Jr. et al. 1985). The presence of open-reef or deep-sea sponges in ‘La Quebrada’ supports the idea of such connections (Gómez-Gutiérrez and Calderón-Gutiérrez 2020). Increased sampling efforts in more mainland anchialine caves can lead to the discovery of new sponge species and increase our knowledge of their biodiversity and biogeography.

With the exception of the sponge B3 *Suberites* sp. n., sponges from Xcalak showed rRNA contigs from non-sponge organisms. *Cinachyrella* sp. n. had contigs associated with polychaetes (Annelida) of the genera *Erinaceusyllis* and *Syllis*, while in *Xestospongia* sp. n., a long contig was assigned to the genus *Diadumene* (Cnidaria). Numerous studies have reported symbiotic relationships between sponges and their associated fauna (Koukouras et al. 1996; Ávila et al. 2007; Wulff 2012; Maldonado et al. 2017; Yu et al. 2020). Polychaetes of the genera *Erinaceusyllis* and *Syllis* have been reported in the anchialine cave of Aerolito on Cozumel Island (Calderón-Gutiérrez et al. 2017). Although the genus *Diadumene* has not been reported in anchialine caves of the Yucatan Peninsula, other organisms from the phylum Cnidaria have been recorded (Calderón-Gutiérrez et al. 2017, 2018). The three-dimensional habitat heterogeneity in sponges, their structural complexity, and their ability to synthesize toxic substances may create environments where other species can live in, or gain an adaptive advantage (Pérez-Botello and Simões 2021). Therefore, sequences of non-sponge organisms found within the sponges suggest a symbiotic relationship, however, this topic requires further exploration.

The tree topologies based on nuclear rRNA and mitochondrial genes were similar and showed that the sponge BE1 and the sponge BE2 were closely related to sponge species belonging to the genera *Cinachyrella* and *Xestospongia* (Demospongiae class). Sponge BE3 was closely related to a marine sponge species of the genus *Suberites* (18S rRNA phylogenetic analysis). Phylogenetic analyses showed *Cinachyrella* sp. n. clustered more closely with *Suberites* sp. n. than to *Xestospongia* sp. n. suggesting closer evolutionary histories between the first two sponges This observation has been previously highlighted in a wide analysis of 68 taxa belonging to Class Demospongiae (Plese et al. 2021).

The total number and order of genes observed in the mt-genomes of *Cinachyrella* sp. n. and *Xestospongia* sp. n. were very similar to those found in other demosponge mt genomes (Wang and Lavrov 2008, Taboada et al. 2018). However, slight differences were observed in the number and arrangement of tRNAs. Identical mitochondrial gene order patterns have been reported at the order level in the class Demospongiae (Plese et al. 2021), and respiratory genes such as nad2-nad5 and atp6-cox3, detected in *Cinachyrella* sp. n. and *Xestospongia* sp. n., have been previously reported as highly conserved (Rosengarten et al. 2008). In this study, analyzed groups belonging to the class Demospongiae (G3–G6) had the same number of rRNAs and in proximity to each other (separated by 1–2 tRNA genes) with the gene order being rrnS-trnG-trnV-rrnL, as reported for several demosponge species (Wang and Lavrov 2008, Taboada et al. 2018).

Future research should focus on describing the biodiversity and phylogenetic relationship of anchialine sponges, using molecular approaches (nuclear and mitochondrial DNA) that lead to better classification and comparison between sponges that can inhabit different environments, such as anchialine caves, seawater (Caribbean sea) and Brackish water.

## Acknowledgments

We would like to thank the Posgrado en ciencias del mar y limnología (PCML-UNAM) and CONAHCYT for their support of this project. We want to thank Efraín Chávez Solís, Erick Sosa Rodríguez, and Luis A. Liévano Beltrán for their invaluable support while navigating the incredible depths and lengths of the anchialine cave system. Our appreciation also goes to Miguel A. Acuapan Acosta for his help with drawings.

## Supplementary tables

**Table S1.** Reference 18S rRNA sequences used for the phylogenetic analysis of the *Cinachyrella* sp. n., *Xestospongia* sp. n., and *Suberites* sp. n.

**Table S2.** Reference mitogenomes used for the phylogenomic analysis and mitochondrial gene arrangements between *Cinachyrella* sp. n., and *Xestospongia* sp. n., demosponge species, and outgroup heteroscleromorph.

**Table S3.** rRNA contigs assembled from metagenomes of the three sponges species (*Cinachyrella* sp. n., *Xestospongia* sp. n., and *Suberites* sp. n.) with %identity, length, coverage and taxonomic identification.

**Table S4.** Comparison of Sponge Species diversity detected in the Anchialine Caves: La Quebrada, El Aerolito, and Xcalak

